# Bradycardia inhibits brain vessel mural cell differentiation via reducing mechanosensory and Jag2-Notch signaling

**DOI:** 10.64898/2026.06.25.734621

**Authors:** Ruchita Shandilya, Sarah J Childs

## Abstract

Bradycardia occurs when the heart rate is lower than normal resulting in reduced cerebral blood flow and contributing to neurodegeneration in adults but how it affects embryonic cerebrovascular development is not well studied. We induce bradycardia by targeting the heart pacemaker channel Hcn4 via chemical (ivabradine) and genetic (*hcn4* mutant) methods. Bradycardia results in reduced brain vessel diameter and mural cell (pericyte and vascular smooth muscle cell) number. Endothelial cells are the first responders in sensing changes in blood flow, and we show that signalling through the canonical endothelial-autonomous mechanosensitive pathway (Piezo1, Mek5, Erk5, Klf2) is reduced in bradycardia. To identify the ligand-receptor combination that transmits signals to developing mural cells, we show that expression of the Notch ligand *jagged2b* is decreased in the brain of both *hcn4* and *klf2* mutants. *jag2b* knockdown reduces mural cell numbers in brain vessels. Restoring *jag2b* levels increases mural cell numbers in both wildtype and *hcn4* mutants. Our work connects bradycardia, mechanosensitive signaling and mural cell recruitment demonstrating that mural cell numbers can be increased in bradycardia by restoring Notch signalling via upregulating endothelial Jag2b.

**Summary:** Bradycardia models show reduced blood flow, Piezo1-*klf2-jag2b*-*notch3* mechanosensing and mural cell recruitment to developing brain vasculature. Restoration of *jag2,* an endogenous endothelial cell ligand, rescues mural cell numbers in bradycardia mutants.

## Introduction

During embryonic brain development, not only blood flow conveys oxygen and nutrients to tissues but also acts as a mechanical cue through providing shear, circumferential and axial stress to the vessel wall. Blood flow shapes development of the endothelial network. Acute or chronic impairments to blood flow to the brain can result in ischemic or hypoxic injury and adverse neurodevelopment (Moreira and Guedes-Martins, 2025). Bradycardia is a condition characterized by slow resting heart rate. Patients present with fatigue, dizziness and syncope owing to reduced cerebral blood flow (Solti et al., 1987). While bradycardia can arise from complex physiological phenomena, it also can have a genetic origin (Ishikawa et al., 2016; Rezazadeh and Duff, 2017). Mutation in sinus node pacemaker channels such as hyperpolarization activated cyclic nucleotide gated potassium channel 4 (HCN4) causes diminished pacemaker currents associated sick sinus syndrome (Wallace et al., 2021). Patients with HCN4 mutations present with reduced heart rate and syncope owing to reduced cerebral blood flow (Laish-Farkash et al., 2010; Milano et al., 2014; Ueda et al., 2004; Wang et al., 2022). HCN4 loss of function mutations have been described in infants and children with bradycardia (Evans et al., 2013; Wacker-Gussmann et al., 2020; Yokoyama et al., 2018). The only intervention available for sinus node dysfunction due to HCN4 channel mutation is pacemaker implantation which significantly improves syncope symptoms, thereby improving the quality of life (Koide et al., 1994; Kusumoto et al., 2019; Rein et al., 1985; Solti et al., 1987). Mouse and fish models with Hcn4 mutations result in bradycardia and reduced pacemaker currents (Baruscotti et al., 2011; Harzheim et al., 2008). Mice with loss-of-function Hcn4 show reduced embryonic brain size (microcephaly), decreased body weight at birth, developmental delays and mortality (Schlusche et al., 2021; Stieber et al., 2003). *hcn4* mutant zebrafish embryos show pericardial edema and reduced heart rate (Liu et al., 2022). While symptoms associated with HCN4 mutation are well studied in heart development and cardiac function, we have little mechanistic information on how bradycardia affects cerebrovascular development.

As the heart and blood vessels are the first organs to form in a developing embryo, blood vessels are exposed to shear stress as soon as they form. Physiological cues from blood flow directs the developing vascular network to adjust (Ghaffari et al., 2015a, b; Jones, 2011a; Udan et al., 2013). Shear force is sensed by mechanosensitive channels and receptors on the luminal side of endothelial cells (ECs). In response to shear forces, endothelial cells undergo changes in their size, morphology and polarity to align in the direction of blood flow (Abe and Berk, 2014; Jones, 2011b; Qu et al., 2023). Diverse flow patterns elicit different responses from ECs. High laminar shear flow inhibits cell cycle progression and stabilizes vasculature, whereas low shear flow or disturbed flow causes high EC turnover (Fang et al., 2017; Nakajima and Mochizuki, 2017). Changes in blood flow patterns are sensed by the mechanosensitive channels present on the luminal side of the endothelium (Baeyens and Schwartz, 2016; Beverley et al., 2025; Davis et al., 2023).

Shear stress sensed by endothelial cells via mechanosensitive channels such as Piezo1, or TRPs increases calcium (Ca^2+^) influx in cells (Santana Nunez et al., 2023; Sonkusare et al., 2012; Thakore and Earley, 2019; Xiao, 2024) that mediates phosphorylation of MEKK3 which in turn phosphorylates downstream Mek5 and Erk5 kinases to drive MEF2-mediated transcription of flow-regulated transcriptional factor, *kruppel-like factor 2 (klf2)* (Campinho et al., 2020), facilitating endothelial cell differentiation, angiogenic sprouting, lumenization and pruning, barrier function, hemodynamic regulation, and vascular homeostasis (Lin et al., 2010; Nayak et al., 2011; Wang et al., 2006; Woo et al., 2008).

Mural cells are an umbrella term for pericytes that wrap smaller vessels like capillaries, and vascular smooth muscle cells supporting large vessels. Mural cells are an integral part of the blood brain barrier (BBB) that stabilize the brain vasculature and fine-tune barrier permeability and vascular tone (Armulik et al., 2010; Nicoli and Grutzendler, 2021; Siekmann, 2023). Disruptions in mural cell numbers or coverage leads to BBB leakage, microvascular aneurysms, and elevated risk of cerebral hemorrhage (Gautam and Yao, 2018; Marbacher et al., 2014; Pan et al., 2021; Sengillo et al., 2013; Sun et al., 2021). Mural cell numbers are impacted substantially in response to changes in the blood flow (Abello et al., 2025; Cheng et al., 2024; Zi et al., 2024). But the mechanism of how changes in blood flow pattern are transmitted to mural cells is incompletely known.

Although it is expressed in endothelial cells, *klf2* plays an integral role in influencing mural cell coverage of blood vessels (Abello et al., 2025; Cheng et al., 2024). CXCL11-CXCR3 (Lee et al., 2024) and CXCL12-CXCR4 (Stratman et al., 2020) promote mural cell chemoattraction via expression of PDGFB downstream of blood flow-induced endothelial *klf2* signaling. Additional non-mechanosensing pathways that communicate between ECs and mural cells in in vitro studies include ANG1-TIE2 (Sundberg et al., 2002; Wakui et al., 2006), PDGF-BB, TGFβ (Hirschi et al., 1998; Qi et al., 2011), , ALK1-BMP (Baeyens et al., 2016) and DLL4- or JAG1-NOTCH3 (Breikaa et al., 2022; Liu et al., 2009; Schulz et al., 2015). Recently, gain of function techniques in zebrafish were used to suggest that Jagged1-Notch3 from endothelium to mural cells is needed for mural cell recruitment during development (Zi et al., 2024). Deletion of Jagged1 from adult endothelial cells results in altered sVMCs in mouse (Breikaa et al., 2022).

Using a physiological model of bradycardia, we show that a reduced heart rate results in reduced blood flow during development, impacting mechanosensing in endothelial cells and initial recruitment of mural cell precursors, affecting the development of brain vasculature. We demonstrate Jag2b is downregulated in bradycardia, but that restoring Jag2b improves mural cell recruitment to brain vessels in developing embryos with bradycardia.

## Results

### Hemodynamic changes downstream of HCN4 loss of function influence brain mural cell numbers

HCN4 is expressed in the heart sinus where it controls heart rate. Loss of HCN4 leads to bradycardia in fish and humans (Fig 1a) (DiFrancesco, 2015; Liu et al., 2022; Nof et al., 2010). As the brain utilizes a large amount of an organism’s blood flow, and perturbations to blood flow affect vascular patterning and mural cell recruitment, we used a bradycardia model to understand how physiological changes to heart rate influence brain vessel development during the sensitive stage when mural cells are being recruited. Embryos expressing the transgenes *Tg(pdgfrβ:gal4:UAS:GFP;kdrl:mCherry)* marking pericytes and endothelial cells were treated with the HCN4 channel blocker ivabradine hydrochloride from 48-75hpf. The timing window was selected to overlap with the period when brain perivascular precursor cells are recruited and differentiate into pericytes (Ando et al., 2019; Bahrami and Childs, 2020). We used changes in heart rate as a proxy for blood flow velocity as these parameters are correlated (Chen et al., 2012; Cheng et al., 2024). We observe a reduction in heart rate with ivabradine treatment, and a decrease in blood flow in the trunk (Movies 1-2). Heart rate is significantly reduced by ∼39% in ivabradine treated embryos with respect to control (Supplementary Figure 1), and there is a significant 37.6% reduction of in brain vessel pericyte numbers (Figure 1A-D), similar to what has been previously reported (Zi et al., 2024). The reduction in heart rate also resulted in a reduction in pericyte process length (Fig 1E, 24.6%) and soma size (Fig 1F, 11.2%). Despite these strong changes to pericytes, vessel diameter remained unchanged (Figure 1G).

**Figure 1:**
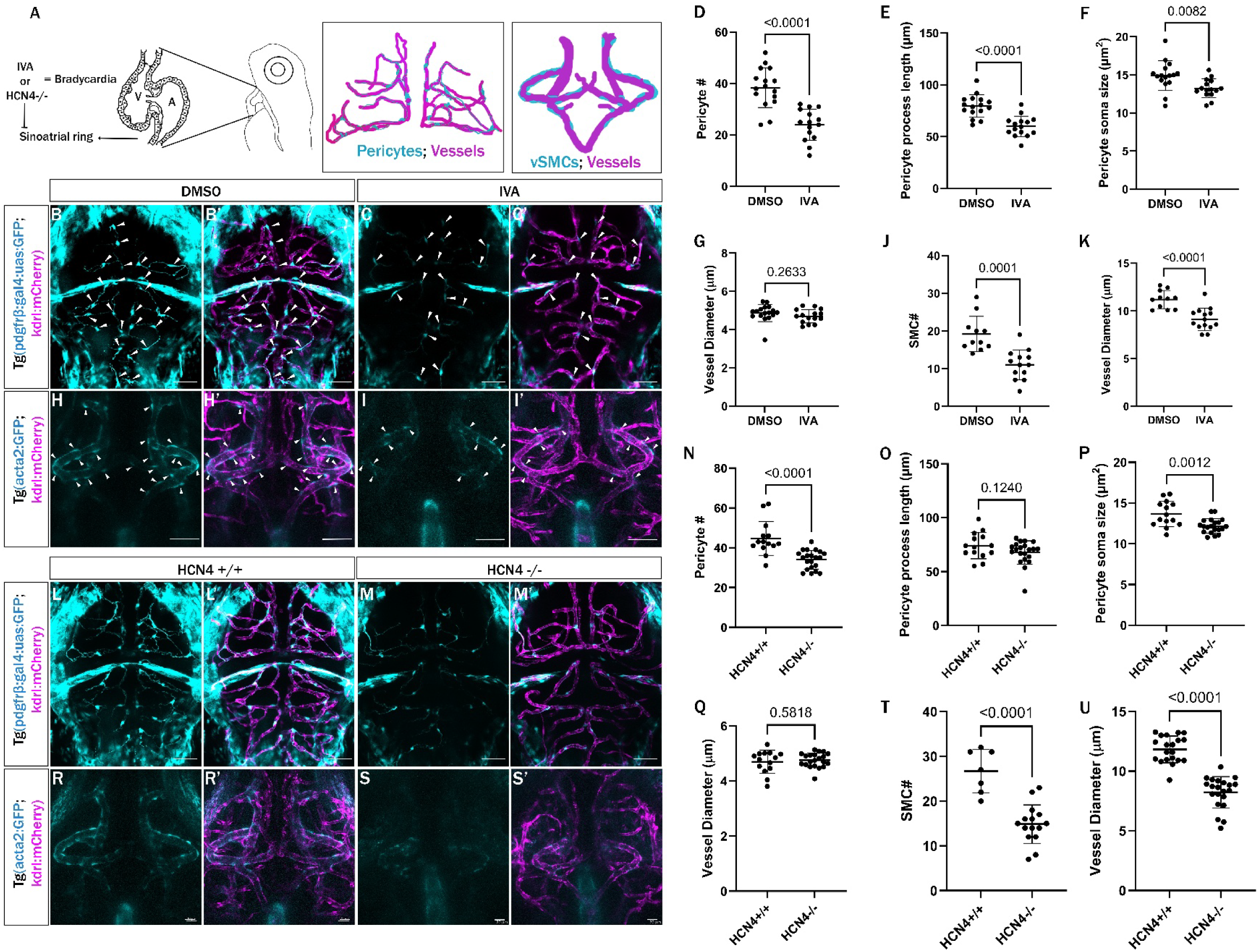
Hemodynamics influences mural cell numbers. A) Schematic illustration shows blood flow reduction using ivabradine hydrochloride (IVA) or genetic HCN4 mutation influences pericytes on brain capillaries and vascular smooth muscle cells (vSMCs) in the Circle of Willis (CoW) arteries. B-C) Confocal images of pericytes and blood vessels in live zebrafish brains of 3 dpf DMSO-treated (B-B’) or 0.3mM IVA-treated (C-C’) *Tg(pdgfrb:GAL4:UAS:GFP; kdrl:mcherry)* zebrafish embryos. Arrows indicate pericytes associated with brain blood vessels. D-G) Quantification of number of pericytes (D), pericyte process length (E), pericyte soma size (F) in the brain vessels and vessel diameter (G) of 3dpf zebrafish embryos treated with DMSO and 0.3mM IVA. (H-I) Confocal images of vascular smooth muscle cells (vSMCs) and blood vessels in the CoW arteries of the brain of DMSO-treated (H-H’) or 0.3mM IVA-treated (I-I’) *Tg(acta2:GFP; kdrl:mcherry)* zebrafish embryos at 4dpf. Arrows indicate the vSMCs associated with CoW of the brain. J) Number of vSMCs in the brain CoW arteries of 4dpf zebrafish embryos treated with DMSO and 0.3mM IVA. K) Vessel diameter of CoW arteries of 4dpf zebrafish embryos. L-M) Pericytes and blood vessels in live zebrafish brains of 3 dpf wildtype (L-L’) and HCN4 maternal zygotic homozygous mutant (M-M’). N-Q) Quantification of pericyte numbers (N), pericyte process length (O), soma size of pericytes (P) and vessel diameter (Q) in the brain vessels of 3dpf wildtype and HCN4 mutant zebrafish embryos. R-S) Vascular smooth muscle cells (vSMCs) and blood vessels in the CoW arteries of the brain of wildtype (R-R’) and HCN4 maternal zygotic homozygous mutant (S-S’) at 4dpf. T) Number of vSMCs in the brain CoW arteries of 4dpf wildtype and HCN4 mutant zebrafish embryos. U) Vessel diameter of 4dpf wildtype and HCN4 mutant zebrafish embryos. Scale bars in images B-I, L-M represent 50µm and images R-S represent 20 µm. Statistics for comparisons use Student’s t-test.

We next counted vascular smooth muscle cells (vSMCs) covering the circle of Willis (CoW) arteries of the brain after Ivabradine treatment on *Tg(acta2:GFP;kdrl:mCherry)* lines from 72-96hpf, a window when precursor cells differentiate into vSMCs (Bahrami and Childs, 2020; Whitesell et al., 2019). With ivabradine treatment, heart rate is significantly reduced by 57.8% (Movie 3, Supplementary Figure 1), and there is a 42.2% reduction in vSMC numbers (Fig 1H-J). The diameter of the CoW arteries is 18.5% reduced (Fig 1K), similar to what has been reported (Cheng et al., 2024)(Supplementary Figure 1, Figure 1H-K).

Using a genetic model of HCN4 maternal zygotic (MZ) mutants, we find a significant reduction in the heart rate of HCN4 MZ mutants by 9.2% and blood flow in trunk vessels at 3 dpf (Movies 4-5, Supplementary figure 1), along with a reduction in pericyte numbers (Fig 1L-N; 23.8%) and pericyte soma size (Fig 1P, 10.9%) with no change in pericyte process length or vessel diameter (Figure 1O, Q). At 4 dpf the heart rate of HCN4 mutant embryos is significantly reduced by 8.2% (Movie 6, Supplementary Figure 1) as were vSMC numbers (Fig 1R-T, 44.3%) and the diameter of the CoW arteries (Fig 1U, reduced 30.4%). This data suggests that bradycardia models significantly reduce recruitment of pericytes and vSMCs, and result in smaller pericyte soma sizes.

### Piezo1 manipulation affects both brain pericyte and vSMC numbers

Piezo1 is a mechanosensitive channel that relays external mechanical forces to intracellular signalling pathways. In the endothelium it relays forces downstream of blood flow. In a late developmental window (3-3.5 dpf and 4-4.5 dpf) activation of Piezo1 using the agonist Yoda1 increases pericyte proliferation and increases brain capillary density (Liu et al., 2020; Zi et al., 2024). To look at effects in an earlier pericyte recruitment window, *Tg(pdgfrβ:gal4:UAS:GFP;kdrl:mCherry)* embryos were treated with Piezo1 50nM Yoda1 agonist or 10nM Dooku1 antagonist from 48-75hpf. We found no significant change in heart rate with either inhibitor (Supplementary figure 1). In this early window, Yoda1 treatment resulted in an increase (33%) and Dooku1 showed a reduction (22.8%) in pericyte numbers with no significant changes in vessel diameter (Supplementary figure 2), similar to previous findings in the later time window. With a 3-4 dpf window of treatment we find an increase in number of vSMCs wrapping around the Circle of Willis with Yoda1 treatment (16.6%) while Dooku1 led to a reduction in vSMC numbers by 28.2% (Supplementary figure 2), similar to how Piezo1 activation and inhibition affects vSMC recruitment in the trunk (Abello et al., 2025). We did not see any changes in the vessel diameter of brain capillaries or Circle of Willis arteries with either treatment (Supplementary figure 2).

### *klf2* expression is reduced in embryos with bradycardia

To test whether bradycardia affects *klf2* expression, we treated embryos with Ivabradine from 48-75hpf and performed HCR-FISH to label *klf2a* mRNA transcripts in the brain. There is a significant reduction (17.2%) in *klf2a* mRNA transcript levels, while expression of the endothelial marker *kdrl* remains unchanged (Figure 2A-2D). A reduction in *klf2a* mRNA levels is also seen in HCN4 MZ homozygous mutants (16.70%), while *kdrl* expression remains unchanged (Figure 2E-H). Thus, reduction of blood flow via HCN4 results in altered expression of *klf2*.

**Figure 2:**
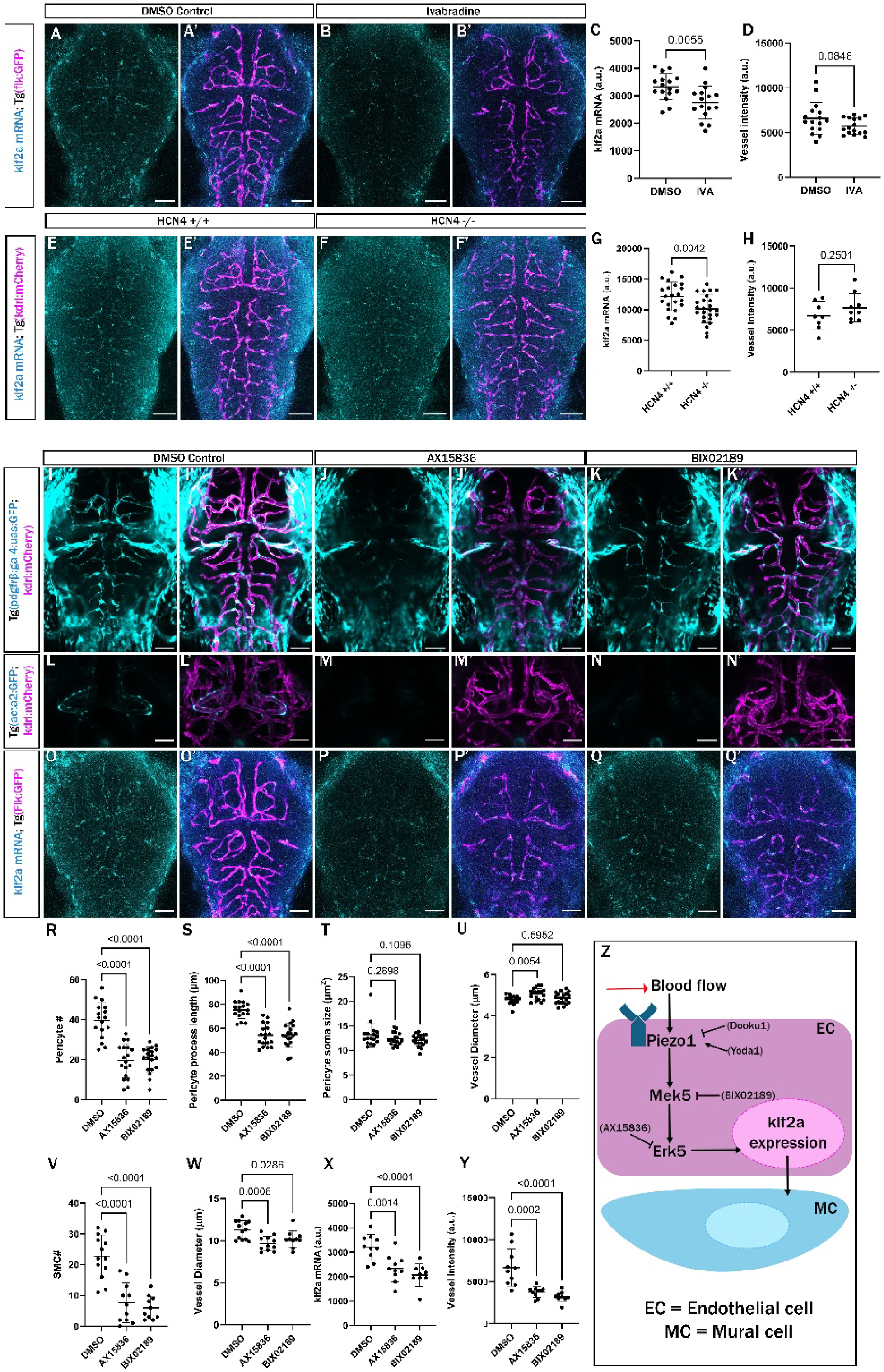
*klf2* influences mural cell numbers via downstream of the Mek5 and Erk5 signalling axis. A-B) Confocal images of the *klf2a* fluorescent in-situ hybridization in the brain blood vessels of DMSO-treated (A-A’) or 0.3mM IVA-treated (B-B’) *Tg(flk:GFP)* zebrafish embryo fixed at 3dpf. C-D) Integrated density of *klf2a* mRNA signal (C) and *Tg(flk:GFP)* signal (D) in the brain blood vessels of 3dpf zebrafish embryos. E-F) *klf2a* fluorescent in-situ hybridization in the brain blood vessels of wildtype (E-E’) or HCN4 MZ homozygous mutant (F-F’) *Tg(kdrl:mCherry)* zebrafish embryo fixed at 3dpf. G-H) Integrated density of *klf2a* mRNA expression (G) and *Tg(kdrl:mCherry)* (H) in the brain blood vessels of 3dpf zebrafish embryos. I-Q) Live confocal images of mural cells and endothelial cells in the brains of embryos labelled with *Tg(pdgfrb:GAL4:UAS:GFP; kdrl:mcherry)* at 3 dpf (I-K), *Tg(acta2:GFP; kdrl:mcherry)* at 4 dpf (L-N) or *Tg(flk:GFP)* and *klf2* HCR ISH at 3 dpf (O-Q) treated with DMSO (I-I’; L-L’; O-O’), 3µM AX15836 (J-J’; M-M’; P-P’) or 3µM BIX02189 (K-K’; N-N’; Q-Q’). Scale bars represent 50µm in all images. R-U) Number of pericytes (R), pericyte process length (S), soma size of pericytes (T) and vessel diameter (U) of the brain vessels of 3dpf zebrafish embryos treated with DMSO, 3µM AX15836 or 3µM BIX02189. V-W) Number of vSMCs (V) and vessel diameter (W) of the brain Circle of Willis arteries of 4dpf zebrafish embryos treated with DMSO, 3µM AX15836 or 3µM BIX02189. X-Y) Integrated density of *klf2a* mRNA signal (X) and *Tg(flk:GFP)* signal (Y) in brain blood vessels of 3dpf zebrafish embryos treated with DMSO, 3µM AX15836 or 3µM BIX02189. Student’s t-tests and one-way ANOVA are used for statistical comparison. Z) Diagram showing the flow of mechanical information via the blood flow through a Piezo1-Mek5-Erk5-*klf2* mechano-signalling axis to influence mural cells.

To understand the molecular mechanism, we examined a pathway identified in vitro where *klf2* expression is controlled downstream of MEKK3 (Zhou et al., 2015; Zhou et al., 2016), in turn, which is downstream of Piezo1 signals and *CamIIK* (Zheng et al., 2022). Inhibiting Mek5 with 3µM BIX02189 or Erk5 with 3µM AX15836 in *Tg(pdgfrβ:gal4:UAS:GFP;kdrl:mCherry)* embryos from 48-75hpf shows no significant changes in the heart rate of AX15836 treated animals but a decrease in heart rate in BIX02189-treated embryos by 11.3% (Supplementary figure 1). We observe a significant reduction in pericyte numbers in both BIX02189 (49.2%) or AX15836 (50.8%) treatment (Figure 2I-2K, 2R-2U). Similarly, there is a severe reduction in the vSMCs in the circle of Willis (66.3% reduction with AX15836 and 73.6% reduction with BIX02189) along with a reduction in vessel diameter by 14.3% with AX15836 and 9.6% with BIX02189 (Figure 2L-2N, 2V-2W). To test whether Mek5 and Erk5 affect *klf2* expression in vivo we used HCR-FISH to measure *klf2a* expression. There is a reduction (27.2% with AX15836 and 35.7% with BIX02189), suggesting that *Mek5* and *Erk5* activity are upstream of *klf2a* expression in vivo (Figure 2O-2Q, 2X). We observed that expression of the *kdrl:GFP* transgene is reduced after either BIX02189 (52.1%) or AX15836 (43.5%) treatment (Figure 2O-2R, 2Y) suggesting additional effects on endothelial cell differentiation. Although different aspects of this pathway have been demonstrated in different cell, mouse and zebrafish models, our complete data assessing both pericyte and vSMC numbers downstream of manipulations of blood flow, Piezo1, Mek5 and Erk5 illustrates an integrated pathway for recruitment of both mural cell types on brain vessels (Figure 2Z).

### *klf2a;klf2b* double genetic mutants shows reduced mural cell numbers

Zebrafish have two KLF2 paralogs, *klf2a* and *klf2b*. Previous work suggests that *klf2a* loss of function is sufficient to alter vSMC recruitment to large trunk veins with no change to the artery (Abello et al., 2025; Cheng et al., 2024; Stratman et al., 2020). However, we find that single *kfl2a* or *klf2b* mutants have no effect on cerebrovascular differentiation. Using a *klf2a* mutant line crossed to reporters *Tg(pdgfrβ:gal4:UAS:GFP;kdrl:mCherry*) at 3 dpf or *Tg(acta2:GFP;kdrl:mCherry)* at 4 dpf we found no effect on pericyte or vSMCs numbers in the CoW or small brain vessels (capillaries and arteries), or vessel diameter (Supplementary Figure 3). Single *klf2b* mutants also show no change in the pericytes or vSMCs at the same time, nor changes on blood vessel diameter (Supplementary Figure 3). However, *klf2a;klf2b* double knockouts on the same backgrounds showed a significant reduction in the heart rate (9.9%), and pericyte number (28.3%) at 3 dpf, without impact on pericyte soma size, pericyte process length or vessel diameter of the brain capillaries (Supplementary figure 1; Figure 3A-B, 3E-3H). Similarly, *klf2a;klf2b* double knockouts at 4 dpf showed reduction of heart rate (13%) and vSMCs (26.2%) (Supplementary figure 1; Figure 3C-D, 3I) without impact on blood vessel diameter of larger arteries (Figure 3J). Thus, loss of the endothelial mechanosensitive transcription factor Klf2 inhibits mural cell recruitment to brain blood vessels.

**Figure 3:**
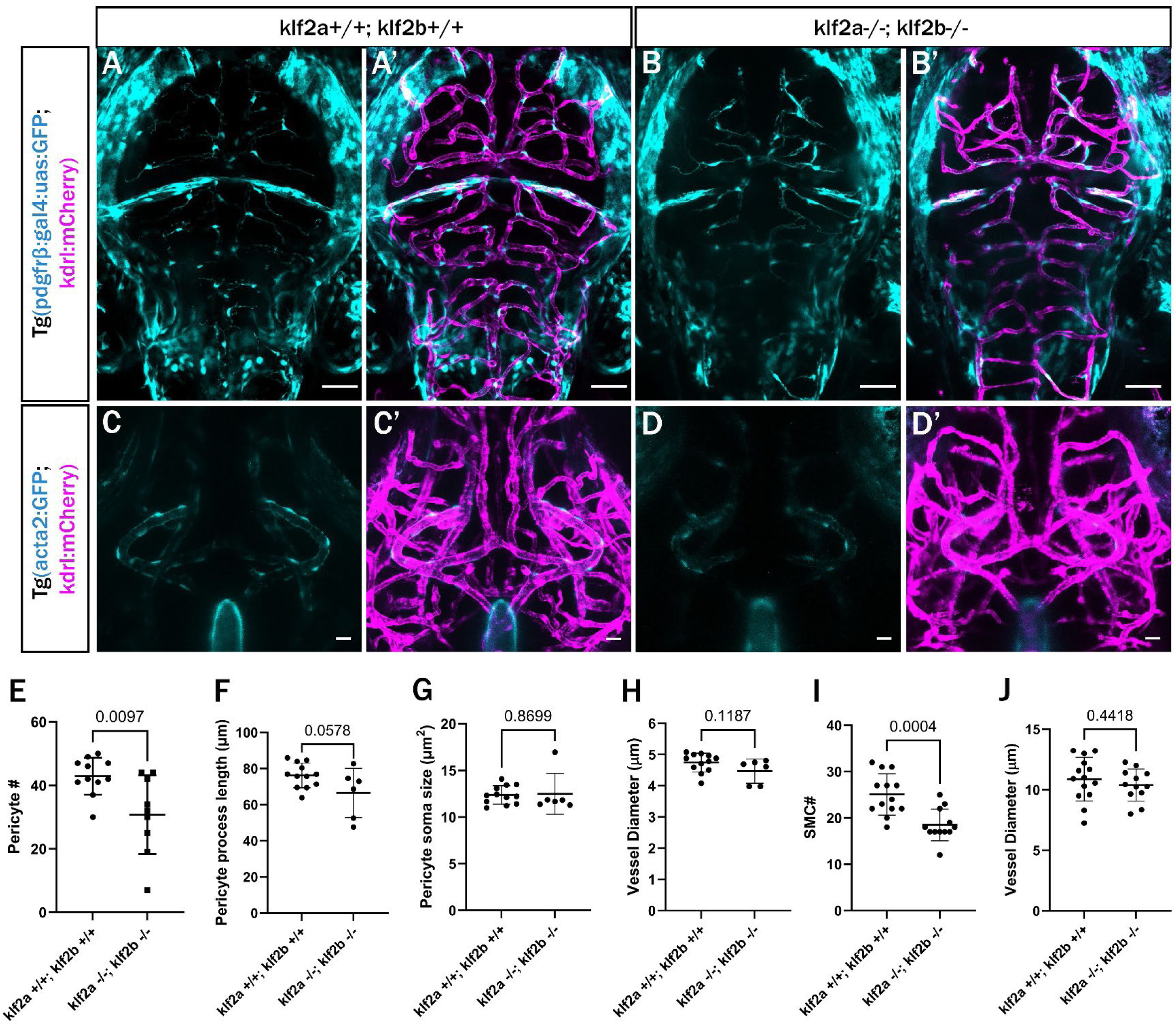
*klf2* double mutants show reduced pericytes and vSMCs. A-) Confocal images of pericytes and blood vessels in live zebrafish brains of 3 dpf wildtype (A-A’) or *klf2a; klf2b* double homozygous mutant (B-B’) *Tg(pdgfrb:GAL4:UAS:GFP; kdrl:mcherry)* zebrafish embryos. C-D) Confocal images of vascular smooth muscle cells (vSMCs) and blood vessels in the CoW arteries of the brain of 3 dpf wildtype (C-C’) or *klf2a; klf2b* double homozygous mutant (D-D’) *Tg(acta2:GFP; kdrl:mcherry)* zebrafish embryos at 4dpf. E-H) Quantification of number of pericytes (E), pericyte process length (F), pericyte soma size (G) in the brain vessels and vessel diameter of brain vessels (H) of 3dpf wildtype or *klf2a; klf2b* double homozygous mutant zebrafish embryos. I-J) Quantification of the number of vSMCs in the brain CoW arteries (G) and vessel diameter (H) of 4dpf wildtype or *klf2a; klf2b* double homozygous mutant zebrafish embryos. Scale bars represent 50µm in images A-B and 20µm in C-D. Statistics for comparisons use Student’s t-test.

### Klf2 influences mural cell numbers via notch signaling

To identify the endogenous signal between endothelial cells and mural cell differentiation we looked to the Notch pathway. Forced expression of Jag1 in zebrafish endothelial cells results in increased brain pericyte coverage (Zi et al., 2024). We investigated if the expression pattern of Notch ligands *dll4*, *jag1*, and *jag2* is changed after manipulating the endothelial mechanosensitive pathway as characterized above. *dll4* expression is reduced with ivabradine treatment but is unchanged after Mek5/Erk5 signaling perturbations (Figure 4A-4D). Likewise, *jag1b* expression is unchanged with ivabradine or Mek5/Erk5 inhibition (Fig 4E-4H). On the other hand, *jag2b* expression was decreased with Ivabradine or AX15836 and BIX02189 treatment (Fig 4I-4L), consistent with being a downstream effector of the pathway in endothelial cells. Single cell sequencing databases from fish and human embryonic/fetal stages show that Jag2 (including Danio rerio *jag2b*) is expressed in brain endothelial cells (Supplementary figure 4). This suggests that Jag2b might be the physiological ligand downstream of Klf2 that interacts with Notch3 receptors on adjacent mural cells (Figure 4M).

**Figure 4:**
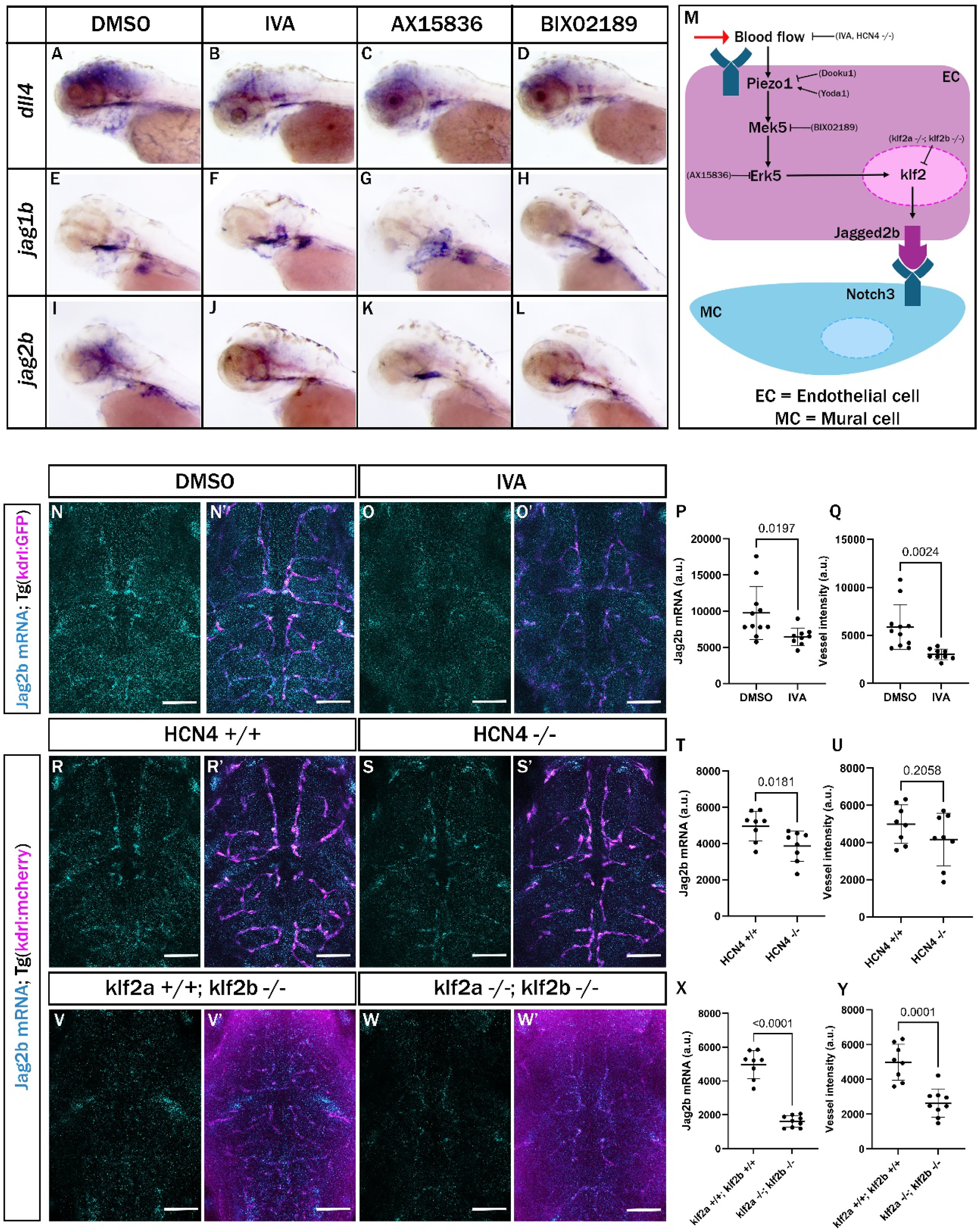
Mechanosensitivity and response to manipulation of Mek5-Erk5 on expression notch ligands. A- L) Lateral view of *dll4* (A-D), *jag1b* (E-H), and *jag2b* (I-L) expression in 3dpf zebrafish embryos treated with DMSO (A, E, I), 0.3mM Ivabradine hydrochloride (B, F, J), 3µM AX15836 (C, G, K) and 3µM BIX02189 (D, H, L) respectively. (N-O) Confocal images of the *jag2b* fluorescent in-situ hybridization in the brain blood vessels of DMSO-treated (N-N’) or 0.3mM IVA-treated (O-O’) *Tg(kdrl:mCherry)* zebrafish embryo fixed at 3dpf. P-Q) Integrated density of *jag2b* mRNA signal (P) and *Tg(kdrl:mCherry)* signal (Q) in the brain blood vessels of 3dpf DMSO or 0.3mM IVA-treated zebrafish embryos. (R-S) Confocal images of the *jag2b* fluorescent in-situ hybridization in the brain blood vessels of wildtype (R-R’) or HCN4 MZ homozygous mutant (S-S’) *Tg(kdrl:mCherry)* zebrafish embryo fixed at 3dpf. T-U) Integrated density of *jag2b* mRNA signal (T) and *Tg(kdrl:mcherry)* signal (U) in the brain blood vessels of 3dpf wildtype or HCN4 MZ homozygous mutant zebrafish embryos. (V-W) Confocal images of the *jag2b* fluorescent in-situ hybridization in the brain blood vessels of wildtype (V-V’) or *klf2a; klf2b* double mutant (W-W’) *Tg(kdrl:mCherry)* zebrafish embryo fixed at 3dpf. X-Y) Integrated density of *jag2b* mRNA signal (X) and *Tg(kdrl:mcherry)* (Y) in the brain blood vessels of 3dpf wildtype or *klf2a; klf2b* double mutant zebrafish embryos. Scale bars are 50µm in all fluorescent in-situ hybridization images. Statistics for comparisons use Student’s t-test.

Reduction in jag2b expression was quantitated using HCR-FISH of *jag2b* in embryos treated with Ivabradine. We find significant reduction in *jag2b* expression (by 33.6%) and *kdrl:GFP* transgene intensity (by 48.4%) in brain blood vessels (Figure 4N-4Q). HCN4 mutants similarly show a decrease in *jag2b* expression (by 22.4%). *kdrl:mCherry* expression is unchanged (Figure 4R-4Y). Similarly, *klf2* double knockouts show reduced *jag2b* mRNA levels (67.7%), and the fluorescence intensity of the *kdrl* transgene was also significantly reduced by 47.1% (Figure 4V-4Y). Changes in *jag2b* expression in multiple models in the mechanosensitive pathway let us to focus on it as a potential endothelial signal to recruit mural cells.

### Jag2b acts as an endogenous mediator of endothelial-mural cell crosstalk downstream of HCN4

Using a validated *jag2b* morpholino (Liu et al., 2007) to knock down its expression, pericyte numbers, pericyte process length and pericyte soma size are significantly reduced by 73.7%, 44.8% and 48.9% respectively, with no effect on vessel pattern or transgene intensity (Figure 5A-5B, 5E-5H). Similarly, there is a significant reduction in vSMCs numbers by 84.2% in the large caliber circle of Willis arteries of the brain (Figure 5C-D, 5I) and a reduction in vessel diameter by 8.2% (Figure 5J). This suggests that *jag2b* is necessary for mural cell recruitment.

**Figure 5:**
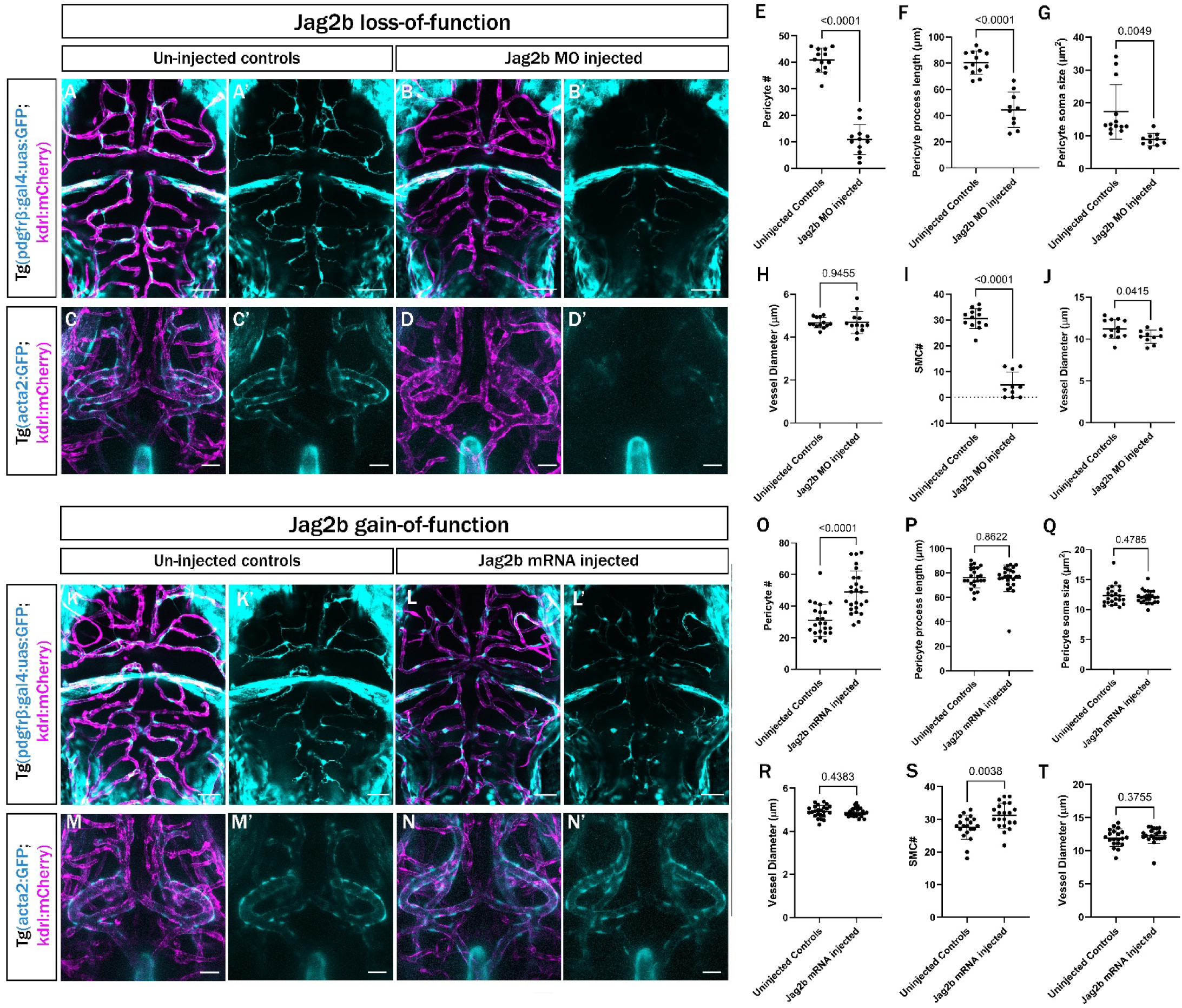
Jag2b loss and gain of function controls mural cell number. A-B) Confocal images of pericytes and blood vessels in live zebrafish brains of 3 dpf uninjected controls (A-A’) or *jag2b* morpholino injected (B-B’) *Tg(pdgfrb:GAL4:UAS:GFP; kdrl:mcherry)* zebrafish embryos. C-D) Vascular smooth muscle cells (vSMCs) and blood vessels in the CoW arteries of the brain of uninjected control (C-C’) or *jag2b* morpholino injected (D-D’) *Tg(acta2:GFP; kdrl:mcherry)* zebrafish embryos at 4dpf. E-H) Quantification of number of pericytes (E), pericyte process length (F), pericyte soma size (G) in the brain vessels and vessel diameter (H) of 3dpf uninjected control or *jag2b* morpholino injected zebrafish embryos. I-J) Quantification of the number of vSMCs in the brain CoW arteries (I) and vessel diameter (J) of 4dpf uninjected control or *jag2b* morpholino injected zebrafish embryos. K-L) Confocal images of pericytes and blood vessels in live zebrafish brains of 3 dpf uninjected controls (K-K’) or *jag2b* mRNA injected (L-L’) *Tg(pdgfrb:GAL4:UAS:GFP; kdrl:mcherry)* zebrafish embryos. M-N) Vascular smooth muscle cells (vSMCs) and blood vessels in the CoW arteries of the brain of uninjected control (M-M’) or *jag2b* mRNA injected (N-N’) *Tg(acta2:GFP; kdrl:mcherry)* zebrafish embryos at 4dpf. O-R) Quantification of pericyte numbers (O), pericyte process length (P), soma size of pericytes (Q) in the brain vessels and vessel diameter (R) of 3dpf uninjected control or *jag2b* mRNA injected zebrafish embryos. S-T) Quantification of the number of vSMCs in the brain CoW arteries (S) and vessel diameter (T) of 4dpf uninjected control or *jag2b* mRNA injected zebrafish embryos. Scale bars in images A-B and K-L represent 50µm and 20µm in C-D and M-N. Statistics for comparisons use Student’s t-test.

To test whether *jag2b* expression is sufficient to increase the number of mural cells, we injected *jag2b* mRNA (Wada et al., 2025) at the single-cell stage in wild type embryos and found that pericyte numbers are increased by 57.7%, while the pericyte process length, soma size and vessel diameter of the brain capillaries are unchanged (Figure 5K-5L, 5O-5R). Similarly, numbers of vSMCs in large vessels of the brain increase by 13.2% with no change vessel diameter (Figure 5M-5N, 5S-5T). This data suggests that increased levels of *jag2b* are sufficient for mural cell recruitment and differentiation process.

Finally, to test whether *jag2b* overexpression rescues the consequences of bradycardia, we injected *jag2b* mRNA in the HCN4 MZ homozygous mutant embryos. *jag2b* gain-of-function completely rescues pericyte numbers (increasing by 126.78%) and process length (increasing by 13.8%) (Figure 6A-6C, 6G-6H). Pericyte soma size and vessel diameter of the brain capillaries are unchanged in both injected embryos and uninjected controls (Figure 6A-6C, 6I-6J). Similarly, vSMCs in large brain vessels are rescued with numbers comparable to the wildtype embryos (an increase by 61.6%) by *jag2b* overexpression (Figure 6D-6F, 6K). Vessel diameter shows an increase of 12.1% (Figure 6D-6F, 6K).

**Figure 6:**
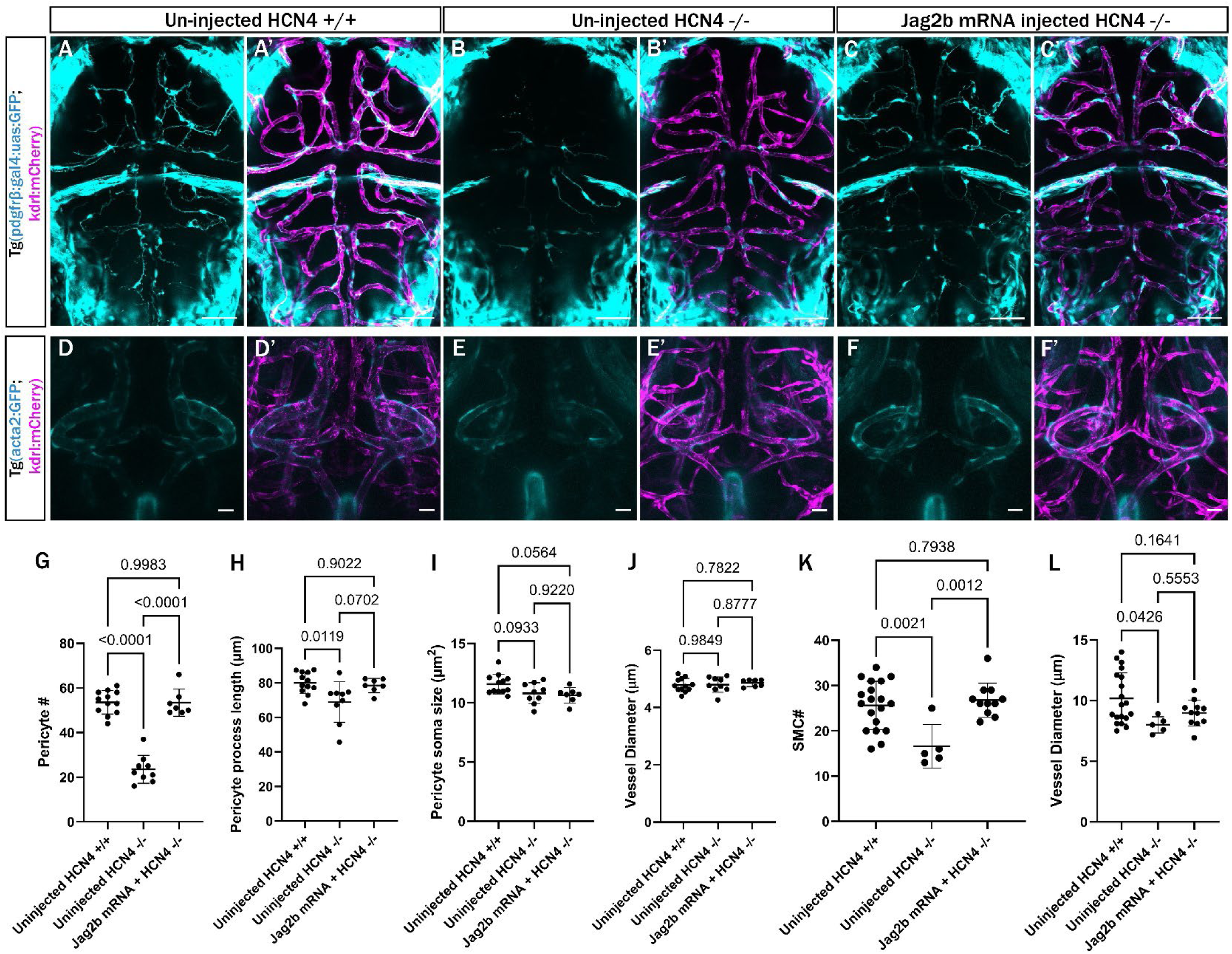
Jag2b overexpression rescues mural cell numbers in HCN4 mutants. A-C) Confocal images of pericytes and blood vessels in live zebrafish brains of 3dpf uninjected wildtype controls (A-A’) or uninjected HCN4 MZ mutant (B-B’) or *jag2b* mRNA injected HCN4 MZ mutant (C-C’) *Tg(pdgfrb:GAL4:UAS:GFP; kdrl:mcherry)* zebrafish embryos. D-F) Vascular smooth muscle cells (vSMCs) and blood vessels in the CoW arteries of the brain of uninjected wildtype controls (D-D’) or uninjected HCN4 MZ mutants (E-E’) or *jag2b* mRNA injected HCN4 MZ mutant (F-F’) *Tg(acta2:GFP; kdrl:mcherry)* zebrafish embryos at 4dpf. G-J) Quantification of number of pericytes (G), pericyte process length (H), pericyte soma size (I) in the brain vessels and vessel diameter (J) of 3dpf uninjected wildtype control or uninjected HCN4 MZ mutants or *jag2b* mRNA injected HCN4 MZ mutant zebrafish embryos. K-L) Quantification of the number of vSMCs in the brain CoW arteries (K) and vessel diameter (L) of 4dpf uninjected wildtype control or uninjected HCN4 MZ mutants or *jag2b* mRNA injected HCN4 MZ mutant zebrafish embryos. Statistics for comparisons use one-way ANOVA.

Taken together, expression of *jag2b* is regulated by upstream mechanosensory pathway members and gain and loss of *jag2b* leads to gain and loss of mural cell recruitment. This suggests that *jag2b* is an endogenous target in mechanosensing mediated endothelial-mural cell communication pathway, facilitating mural cell recruitment and differentiation on brain blood vessels.

## Discussion

This study investigates the mechanism of how external mechanical forces from blood flow are transmitted by endothelial cells to regulate the communication between endothelial and mural cells, influencing mural cell recruitment and differentiation to achieve a stabilized vasculature. We started with a model of bradycardia, using HCN4 mutants. There are many different mechanisms by which patients have bradycardia, but some patients with sick sinus syndrome carry HCN4 mutations and experience syncope owing to reduced cerebral blood flow, often requiring a pacemaker intervention (Koide et al., 1994; Ueda et al., 2004; Wang et al., 2022). We find that maternal-zygotic, but not zygotic *hcn4* mutants show a mural cell phenotype, suggesting that the *hcn4* transcript is maternally deposited. There is a Hcn4-like (*hcn4l*) gene in zebrafish which may partially compensate for loss of *hcn4* (Liu et al., 2022; von der Heyde et al., 2020), although our *hcn4* mutants show a phenotype without losing *hcn4l*. In contrast, Ivabradine inhibits both Hcn4 and Hcn4l channels. Hcn4 mutants have been previously characterized in zebrafish only in terms of heart function. The role of bradycardia in *hcn4* mutants in cerebrovascular development is not well-explored.

Piezo1-mediated mechanosensing alters *klf2a* expression and mural cell numbers (Abello et al., 2025; Zheng et al., 2022). Several of the endothelial mechanosensing pathway components have been identified in cell culture but their in vivo roles have not been tested. We inhibited Mek5 and Erk5 via small molecule inhibitors and find that both strongly modulate *klf2* expression and mural cell numbers. This suggests that Mek5 and Erk5 communicate blood flow associated mechanical information to *klf2* as the central mechanosensitive transcription factor in vivo. We observe a change in vessel diameter and total vessel network with Mek5 and Erk5 inhibition potentially due to the crucial role of Mek5 and Erk5 in cardiovascular development. Vessel enlargement after the Mek5 and Erk5 inhibition may be due to reduced mural cell numbers. Mek5 and Erk5 inhibition reduce both *klf2a* and *klf2b*, affecting the mechanosensing to mural cells. Neither *klf2a* nor *kl2b* single mutants displayed any changes in mural cell numbers consistent with previous findings, (Cheng et al., 2024; Novodvorsky et al., 2015; Rasouli et al., 2018) but *klf2a; klf2b* double knockout mutants exhibited a significant decrease in both pericytes and smooth muscle cells in brain blood vessels.

To narrow down which endothelial ligands might communicate to mural cells downstream of the mechanosensing pathway we used multiple upstream interventions (reducing blood flow-and Mek5/Erk5-inhibition) coupled with an expression screen of candidate ligands. The use of multiple upstream manipulations allowed us to focus on genes that were affected by all the manipulations. Of the possible Notch ligands, we found *jag2b* expression is influenced by flow and Mek5-Erl5 inhibition and its expression is enriched in endothelial cells, while its receptor, Notch3 is expressed in mural cells across human, mouse and zebrafish. Notch ligands Jag1 and DLL4 are involved in vascular development, mural cell recruitment and differentiation and maintenance of the barrier integrity of the vessels (Boscolo et al., 2011; High et al., 2008; Liu et al., 2009), but Jag2 has not been previously analyzed in this context. Notch3 is essential for regulating pericyte recruitment and differentiation process; and Notch3 mutants have reduced pericytes and vSMCs, contributing to pathologies such as vascular cognitive impairment, cerebral small vessel diseases, stroke and dementia (Liu et al., 2009; Liu et al., 2010; Wang et al., 2014).

We showed that *jag2b* expression in endothelial cells is significantly reduced in brain vessels of multiple models of mechanosensing pathway models (bradycardia, MEK5/ERK5 pathway and *klf2* mutants). *Jag2b* knockdown not only reduced mural cell numbers but also influenced their phenotype, in terms of soma size and process length. *jag2b* overexpression rescued this reduction. suggesting that Jag2b and Notch3 may interact to control mural cell numbers and phenotype in the brain vasculature. Our finding outlines a flow-Mek5-Erk5-klf2-jag2b-notch3 mechanosignalling axis with *jag2b* acting as a key transmitter molecule that relay blood flow-*klf2* mediated mechanical information from endothelial cells to mural cells, regulating their recruitment and differentiation. In developing fish and humans, jag2 is enriched in endothelial cells.

While pacemakers are mostly implanted postnatally for arrythmias, our data suggests that bradycardia during developmental stages could have profound effects on the developing cerebrovasculature. Pacemaker intervention may become critical during fetal development, as prolonged bradycardia in utero impairs cerebral perfusion and contributes to cerebrovascular pathologies. Significant pre-clinical research has been made to develop a minimally invasive fetal micro-pacemakers or percutaneously installed pacemakers using surgical intervention (Bar-Cohen et al., 2015; Zhou et al., 2014). However, there is still a long way to go to implement these novel therapeutic approaches in humans. The rescue experiments in our work demonstrate that cerebrovascular pathologies owing to bradycardia could be managed if mechanosensing could be restored to normal levels. This information could help in developing a potential non-invasive therapeutic approach for managing proper brain vascular development in the presence of fetal bradycardia, thereby improving the quality of life postnatally.

## Materials and methods

### Zebrafish husbandry and strains

All experimental investigations were performed in compliance with the Canadian Council on Animal Care, and ethics approval was granted by the University of Calgary Animal Care Committee (AC25- 0151). *Danio rerio* embryos were maintained in E3 medium at 28°C (Westerfield, 1995). E3 water was replaced with 1X PTU (N-Phenylthiourea - Sigma-P7629) to prevent pigment development. Sex is not taken into consideration in experiments as zebrafish sex is not determined until 28dpf. Transgenic lines used in the present work include: *TgBAC(pdgfrβ:GAL4FF)^ca42^*(Whitesell et al., 2019)*, TgBAC(4xUAS:EGFP)^mpn100Tg^*(DeMaria et al., 2013)*, Tg(acta2:GFP)^ca7^*(Whitesell et al., 2014)*, Tg(kdrl:mCherry)^ci5^*(Proulx et al., 2010)*, Tg(kdrl:GFP)^la116^*(Choi et al., 2007). Zebrafish lines with a point mutation (T>A) in allele of *hcn4* (*Tg(hcn4^sa11188^*); point mutation (A>T) in allele of *klf2a* (*Tg(klf2a^sa24120^*) and point mutation (C>A) in allele of *klf2b* (*Tg(klf2b^sa43252^*) were obtained from the Zebrafish International Resource Center (ZIRC). All the embryos from heterozygous intercrosses were subjected to blinded quantification prior to genotyping.

### Genotyping

Adult fish genotyping was performed by anesthetizing fish in 0.016% of tricaine and clipping a small part of the tailfin. For embryo and larval genotyping, whole embryos were anesthetized with 0.4% tricaine and sampled. The hotshot protocol was used for the gnomic DNA (gDNA) isolation by incubation in Base solution (50mM NaOH) at 95°C for 30 minutes. Post-incubation, Neutralization solution (1M Tris-HCl pH 8) was added.

Custom TaqMan™ SNP Genotyping Assays were acquired for *hcn4^sa11188^* (ANZTYH2), *klf2a^sa2412^*^0^ (ANRWZFR) *klf2b^sa43252^* (ANYM7GY; ThermoFisher Scientific) and performed as per the manufacturer’s instructions using TaqMan™ Fast Advanced Master Mix (ThermoFisher – 4444557) and custom TaqMan™ SNP Genotyping Assay probes (20X) specific for the target genes and isolated gDNA before PCR and analysis on the QuantStudio 6 Flex Real-Time PCR System (Applied Biosystems). The wildtype and mutant allele are reported in VIC and FAM, respectively.

### Drug treatments

Prior to treatments, embryos were dechorionated and maintained in 1X PTU (N-Phenylthiourea) at 28°C in 24-well plates with ∼10-15 embryos per well. Stock solutions of all small-molecule inhibitors were prepared in DMSO. 0.3mM Ivabradine hydrochloride (Sigma SML0281) targets Hcn4. 50nM Yoda1 is a Piezo1 agonist (Sigma SML1558) and 10nM Dooku1 is a Piezo1 antagonist (Sigma SML2397). Mek5 was inhibited using BIX02189 (Sigma SML2355) and Erk5 by AX15836 (Tocris Bioscience 5843/10) (Drew et al., 2012; Lochhead et al., 2020; Tatake et al., 2008). Control media used equal amounts of DMSO (0.1%).

### Heart rate measurements

For heart rate measurements, embryos were mounted in low-melt agarose without tricaine and observed using the MicroZebraLab apparatus (Viewpoint Life Sciences - ViewPoint Behaviour Technology) at 10x magnification. Heart rate was manually counted over 30s from movies.

### Confocal microscopy

Zebrafish were anesthetized using 0.4% tricaine prior to mounting on glass-bottomed petri dishes (MatTek Corp - P50G-1.5-30-F) using 0.8% low-melt agarose. A Zeiss LSM900 microscope was used for confocal imaging with a 20 X (0.8 NA) objective. Zen Blue and Fiji (Schindelin et al., 2012) software were used for image analysis post-imaging.

### Vascular quantification

Pericyte numbers in the brain vasculature were manually counted using Fiji counting tool in flattened Z-stacks of the brain vasculature of 3dpf zebrafish embryos. For process length, individual process on either side of the pericyte soma were manually measured and averaged to get the total pericyte process length (µm) in an embryo. To measure pericyte soma size, individual soma boundaries were traced manually and averaged to get total pericyte soma size (µm^2^) in an embryo. For the vSMC counts, individual soma along the circle of Willis arteries were counted. All the data were scored blinded to genotype.

Vessel diameter and total vessel network length was measured using Python-based VesselMetrics software using the confocal images of 3dpf embryos in all treatments and across all genotypes (McGarry et al., 2024).

### In-situ hybridization

Embryos were fixed in 4% paraformaldehyde, stored in methanol at −20, then permeabilized with proteinase K (1 mg/mL stock; Invitrogen, 4333793) at concentrations depending on the developmental age. After permeabilization, embryos were pre-hybridized and hybridized with 50% formamide hybridization buffer (50% formamide, 5X SSC, 5mg/mL torula yeast RNA, 50µg/mL heparin, 0.1% tween-20, 10x dextran sulfate) and probes for the target genes (*dll4, jag1b* and *jag2b*) at 60°C. After probe hybridization, embryos were washed twice with wash buffer (50% formamide, 2X SSC and 0.1% Tween-20) for 5 minutes, once with 2X SSC for 15 minutes and twice with 0.2X SSC for 30 minutes at 60°C. After washing, the embryos were subjected to blocking with 8% non-specific sheep serum (NSS) in PBT for an hour. Post-blocking embryos were bound with anti-digoxigenin F_AB_ fragments conjugated with alkaline phosphatase in 1% NSS/PBT. Probes were detected using NBT/BCIP diluted in NTMT buffer (100mM Tris-HCl, 100mM NaCl, 100mM MgCl_2_, 0.1% Tween 20) and fixed with 4% PFA followed by addition of glycerol overnight prior to imaging.

For a quantitative analysis of mRNA expression, Hybridization Chain Reaction (HCR v3.0, Molecular Instruments) was performed using custom probes for *klf2a, kdrl* and *jag2b* as per the manufacturer’s instructions. All data were blinded for analysis.

### Integrated density analysis

*klf2a*, *jag2b* and *kdrl* intensity in 3dpf fluorescent in-situ embryos was analyzed using ImageJ software. The freehand draw tool was used to mark a region of interest and used to measure the overlapping *klf2a* or *jag2b* expression. For corrected mean integrated density, mean background intensity was subtracted from the mean intensity of mRNA or vessel and the total intensities in each embryo were averaged.

### Microinjections

*jag2b* morpholino (5’-TCCTGATACAATTCCACATGCCGCC-3’) (Jacobs et al., 2022; Liu et al., 2007) has been published. *jag2b* plasmid cDNA was a kind gift from Dr. Isao Kobayashi, Institute of Science and Engineering, Kanazawa University, Ishikawa, Japan (Wada et al., 2025). *jag2b* cDNA with pCS2+ was linearized with and mRNA prepared using Invitrogen mMESSAGE mMACHINE SP6 (AM1340) kit. Embryos from a wildtype or *hcn4* homozygote in-cross were injected with *jag2b* morpholino or *jag2b* mRNA at a single-cell stage.

### Statistical analysis

All inhibitor treated groups were compared to DMSO control groups. All mutants were statistically compared to sibling wildtypes. GraphPad prism-10.6.1 software was used for statistical analysis, and significance was determined by p<0.05. The tests used for statistical analysis are mentioned in figure legends. For the comparison of two groups, a two-tailed unpaired t-test was used while one-way ANOVA was used for multi-groups analysis. All results on the graphs represent mean ± standard deviation.

## Acknowledgements

We thank Danielle Blackwell and Sonia Aksamit for zebrafish care. We are grateful to Childs lab members for encouragement and feedback and Dr. Jae-Ryeon Ryu for technical expertise. The *jag2b* morpholino was a kind gift from Dr. Peng Huang. We acknowledge the Alberta Children’s Hospital Research Institute Imaging core for microscopes and technical assistance.

## Data and resources availability

All details of resources are provided in the supplementary information.

## Author contributions

Conceptualization-: SJC; Investigation: RS, SJC; Methodology: RS, SJC; Resources: SJC; Writing- original draft: RS, SJC; Writing- reviewing and editing- RS, SJC

## Supplemental data

**Supplemental Figure 1:**
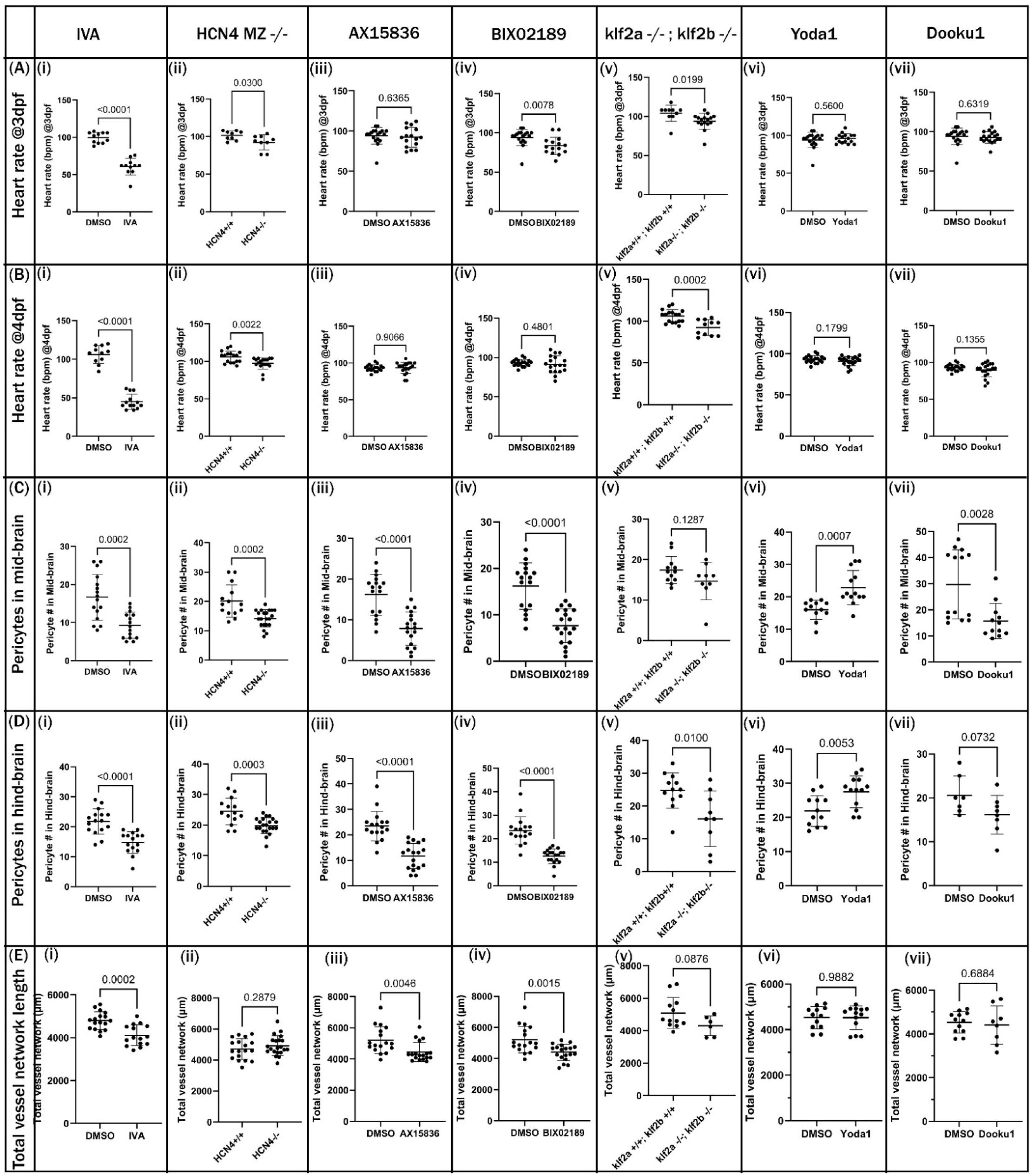
Effect of small-molecule inhibitors and corroborative mutation on hemodynamics and pericytes in midbrain and hindbrain and total vessel network. A-E) Graphs represent heart beats per minute (bpm) at 3dpf (A) and 4dpf (B); pericyte numbers in midbrain (C) and hindbrain (D) and total vessel network length (E) in DMSO, 0.3mM IVA (i), HCN4 mutants (ii), 3µM AX15836 (iii), 3µM BIX02189 (iv), klf2 double mutant (v), 50nM Yoda1 (vi) and 10nM Dooku1 (vii) treated zebrafish embryos. Statistics use Student’s t-test.

**Supplemental figure 2:**
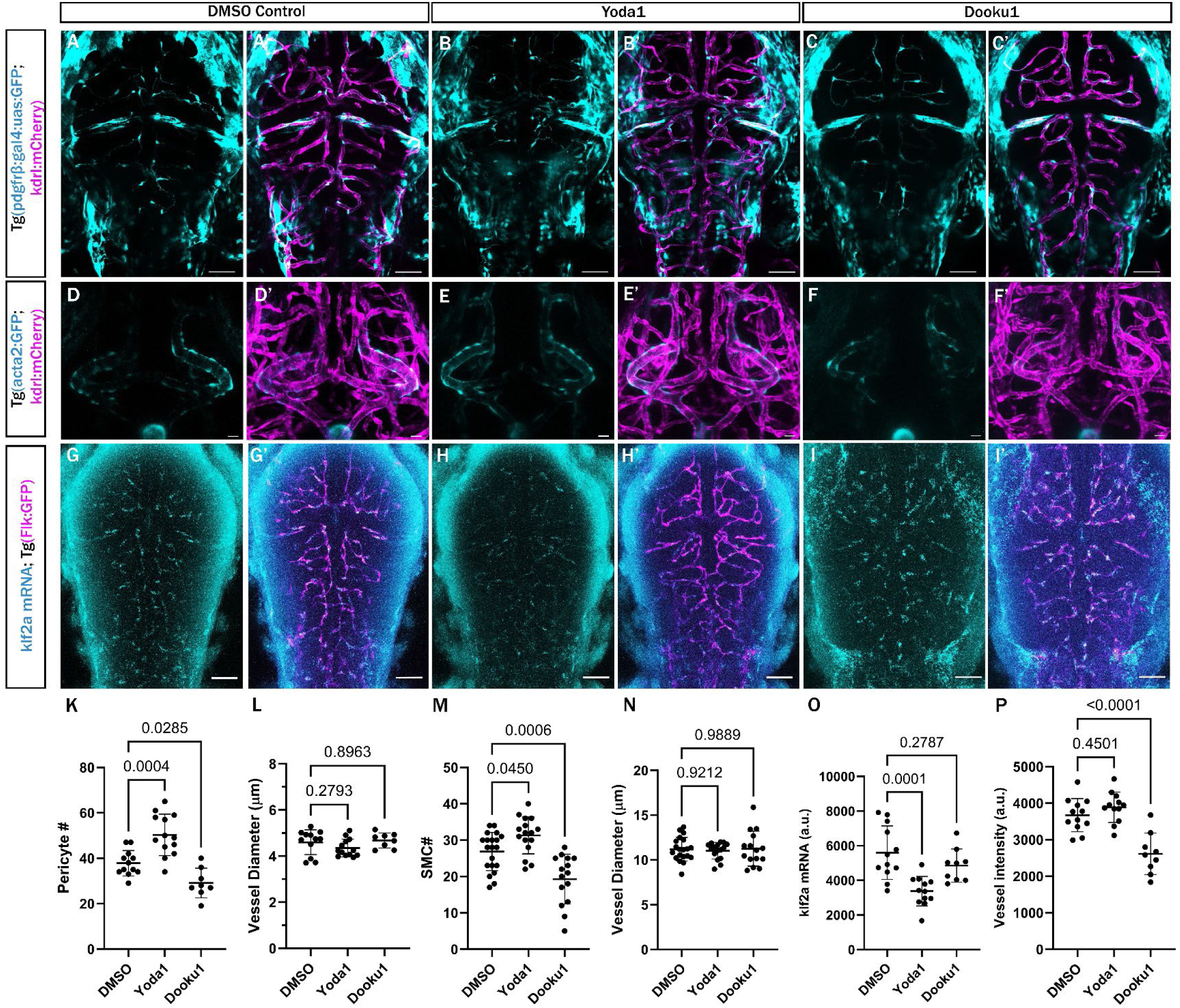
Mechano-sensing modulation via Piezo1 affects mural cell numbers and expression of *klf2.* A-I) Confocal live images of mural cells and endothelial cells in the brains of embryos labelled with *Tg(pdgfrb:GAL4:UAS:GFP; kdrl:mcherry)* at 3 dpf (A-C), *Tg(acta2:GFP; kdrl:mcherry)* at 4 dpf (D-F) or *Tg(flk:GFP)* and klf2 HCR-ISH at 3 dpf (G-I) treated with DMSO (A-A’; D-D’; G-G’), 50nM Yoda1 (B-B’; E-E’; H-H’) or 10nM Dooku1 (C-C’; F-F’; I-I’). Scale bars are 50µm in A-C and G-I; 20µm in D-F. K-L) Number of pericytes (K) and diameter of brain vessels (L) of 3dpf zebrafish embryos treated with DMSO, 50nM Yoda1 and 10nM Dooku1. M-N) Number of vSMCs (M) and diameter of the brain CoW arteries (N) of 4dpf zebrafish embryos treated with DMSO, 50nM Yoda1 and 10nM Dooku1. O-P) Integrated density of *klf2a* mRNA signal in brain vessels (O) and *Tg(flk:GFP)* signal intensity of brain blood vessels (P) of 3dpf zebrafish embryos treated with DMSO, 50nM Yoda1 and 10nM Dooku1. Statistics use Student’s t-test.

**Supplemental figure 3:**
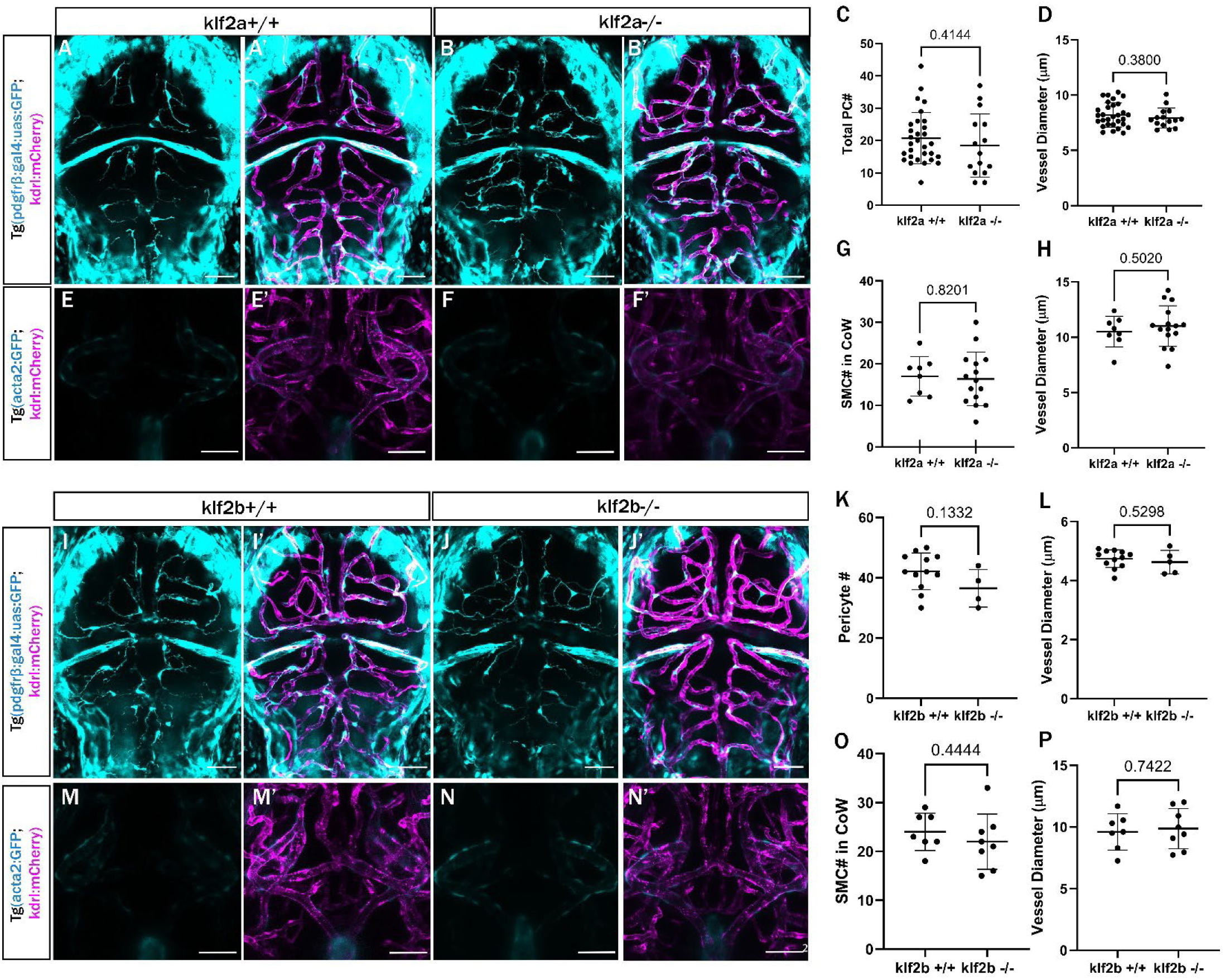
Single mutation of *klf2a* or *klf2b* have no effect of mural cell numbers. A-B) Confocal images of pericytes and blood vessels in live zebrafish brains of 3 dpf wildtype (A-A’) or *klf2a* homozygous mutant (B-B’) *Tg(pdgfrb:GAL4:UAS:GFP; kdrl:mcherry)* zebrafish embryos. C-D) Quantification of number of pericytes in the brain vessels (C) and vessel diameter (D) of 3dpf wildtype or *klf2a* homozygous mutant zebrafish embryos. E-F) Confocal images of vascular smooth muscle cells (vSMCs) and blood vessels in the CoW arteries of the brain of 3 dpf wildtype (E-E’) or *klf2a* homozygous mutant (F-F’) *Tg(acta2:GFP; kdrl:mcherry)* zebrafish embryos at 4dpf. G-H) Quantification of the number of brain vSMCs (G) and vessel diameter (H) of the brain CoW arteries 4dpf wildtype or *klf2a* homozygous mutant zebrafish embryos. I-J) Confocal images of pericytes and blood vessels in live zebrafish brains of 3 dpf wildtype (I-I’) or *klf2b* homozygous mutant (J-J’) *Tg(pdgfrb:GAL4:UAS:GFP; kdrl:mcherry)* zebrafish embryos. K-L) Quantification of number of pericytes in the brain vessels (K) and vessel diameter (L) of 3dpf wildtype or *klf2b* homozygous mutant zebrafish embryos. M-M) Confocal images of brain vascular smooth muscle cells (vSMCs) and blood vessels in the CoW arteries of the brain of 3 dpf wildtype (M-M’) or *klf2b* homozygous mutant (N-N’) *Tg(acta2:GFP; kdrl:mcherry)* zebrafish embryos at 4dpf. O-P) Quantification of the number of vSMCs in the brain CoW arteries (O) and vessel diameter (P) of 4dpf wildtype or *klf2b* homozygous mutant zebrafish embryos. Scale bars represent 50µm in all images. Statistics for comparisons use Student’s t-test.

**Supplementary figure 4:**
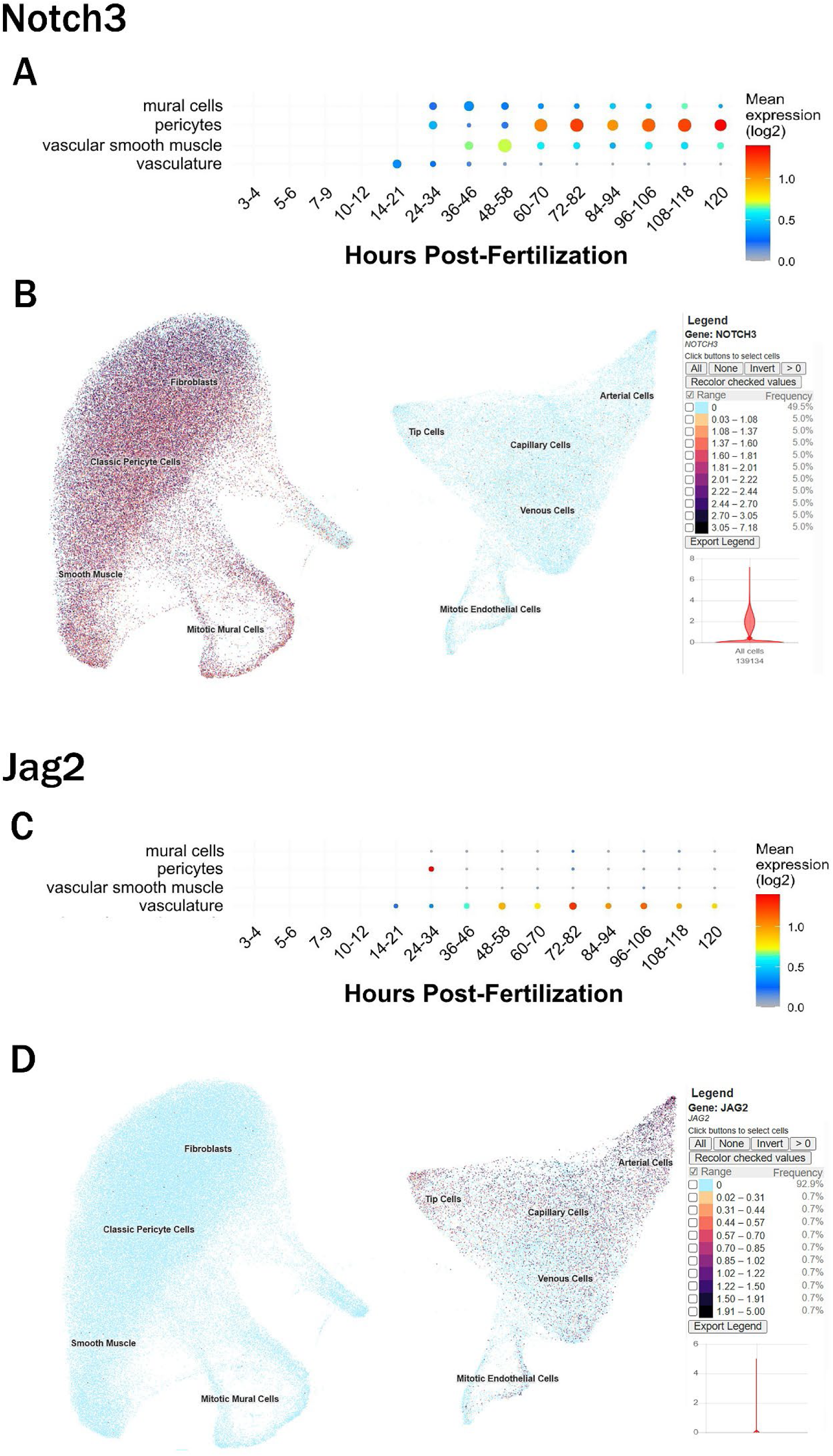
Database showing the enrichment of Notch3 in mural cells and Jag2 in endothelial cells in fish and human. Expression of *notch3* in (A) single cell-RNAseq analysis at different developmental stages in endothelial and mural cells (Sur et al., 2023); (B) Single cell analysis of the brain in human fetal development (Speir et al., 2021); (C) Expression of *jag2b* in zebrafish at different developmental time-points (Sur et al., 2023); (D) single cell-RNAseq analysis showing jag2 expression in different cell clusters of the human fetal brain (Speir et al., 2021).

### Key resources table

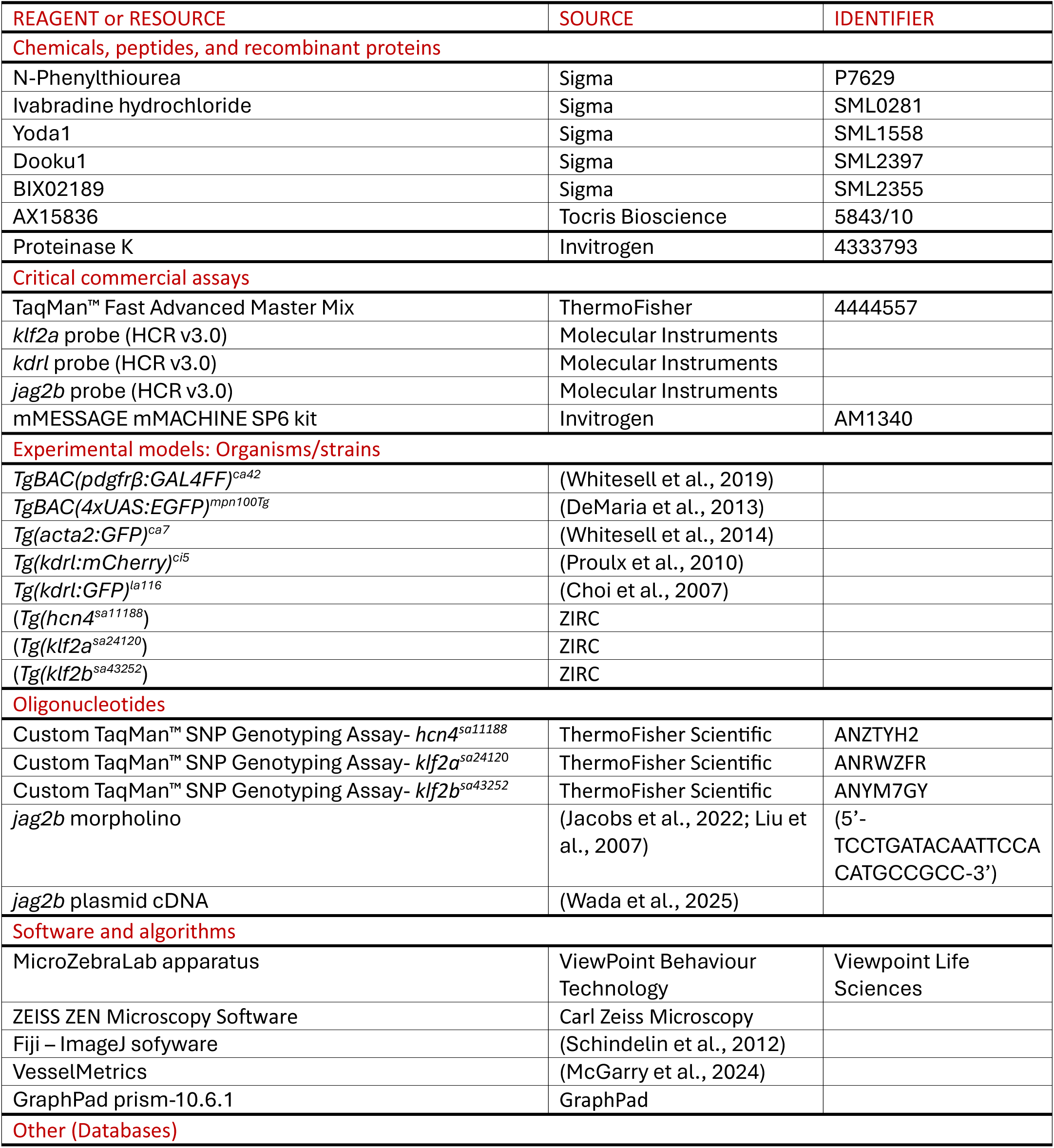
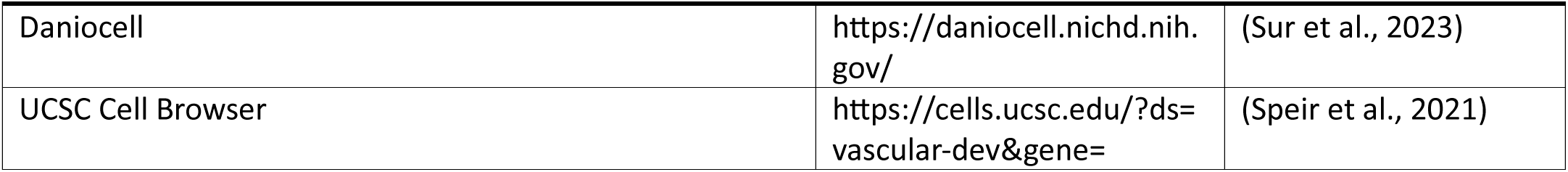

### Movie Captions

**Movie 1.** Heart rate in 3dpf DMSO vs Ivabradine treated zebrafish embryo

**Movie 2**. Blood flow in trunk vessel (dorsal aorta) of 3dpf DMSO vs Ivabradine treated zebrafish embryo

**Movie 3.** Heart rate in 4dpf DMSO vs Ivabradine treated zebrafish embryo

**Movie 4.** Heart rate in 3dpf wildtype vs HCN4 MZ mutant zebrafish embryo

**Movie 5**. Blood flow in trunk vessel (dorsal aorta) of 3dpf wildtype vs HCN4 MZ mutant zebrafish embryo

**Movie 6.** Heart rate in 4dpf wildtype vs HCN4 MZ mutant zebrafish embryo

